# PhyGraFT: a network-based method for phylogenetic trait analysis

**DOI:** 10.1101/2022.05.29.493939

**Authors:** Hirotaka Matsumoto, Motomu Matsui

## Abstract

With the determination of numerous viral and bacterial genome sequences, phylogeny-trait associations are now being studied. In these studies, phylogenetic trees were first reconstructed, and trait data were analyzed based on the reconstructed tree. However, in some cases, such as fast evolution sequences and gene-sharing network data, reconstructing the phylogenetic tree is challenging. In such cases, network-thinking, instead of tree-thinking, is gaining attention. Here, we propose a novel network-thinking approach, PhyGraFT, to analyze trait data from the network. We validated that PhyGraFT can find phylogenetic signals and associations of traits with the simulation dataset. We applied PhyGraFT for influenza type A and virome gene-sharing datasets. As a result, we identified several evolutionary structures and their associated traits. Our approach is expected to provide novel insights into network-thinking not only for typical phylogenetics but also for various biological data, such as antibody evolution.

## Introduction

With advancements in DNA-sequencing technologies, accumulated genome sequences and various trait data shed light on the phylogeny-trait associations of numerous microbial organisms (1) and viruses (2). In previous studies, phylogenetic methods play an essential role in elucidating the relationships between evolutionary structure and trait data. For example, the Parsimony score (3) and association index (4) are frequently used to evaluate evolutionary associations of discrete traits based on a given phylogenetic tree. Other phylogenetic methods and tools for analyzing continuous traits (5, 6), phylogenetic uncertainty (7), and the state-transition process (e.g., BiSSE) (8) have been developed to analyze phylogeny-trait associations.

In virology, phylogeny-trait associations, such as virulence (9), population structure and geographic location (10), host tissues (11), and host cell types (12) have been studied (13). However, phylogenetic trait analysis of viruses is still challenging due to their rapid evolution in general. For example, influenza virus envelope proteins accumulate mutations rapidly due to antigenic drift. Moreover, antigenic shift with reassortment further complicates the evolutionary history. Additionally, there are other cases in which we can quantify similarities but cannot convert these into evolutionary distances, such as gene-sharing networks (14, 15) or protein structure similarity networks (16). Such networks appear not only for studying typical evolution but also for various research into bacterial evolution, including horizontal gene transfer (17) and antibody evolution (18). In such cases, the phylogenetic tree-based methods cannot be used. To overcome these difficulties, network-thinking approaches, instead of tree-thinking approaches, have been proposed as novel methods for phylogenetic analyses (19).

In graph and network science, graph Laplacian is widely used, since the eigenvectors of graph Laplacian are known to represent the basis of the graph. For example, the eigenvectors of small eigenvalues are used for the low dimensional representation of the graph, namely graph embedding (20). The graph Laplacian is also used in phylogenetic research. For example, the eigenvalues of the graph Laplacian derived from the phylogenetic distance matrices are used for classifying phylogenetic trees (21, 22). This approach has also been extended to analyze phylogenetic trait evolution by constructing a graph Laplacian, considering both evolutionary and trait similarity (23). Moreover, a novel phylogenetic tree reconstruction method based on the graph Laplacian of sequence similarity was proposed (24). This uses the eigenvector of the second smallest eigenvalue, called the Fielder vector, for top-down tree reconstruction. Therefore, the network-based method using graph Laplacian is a promising approach for novel phylogenetic analyses.

In this research, we proposed a novel network-based phylogenetic method, Phylogenetic method with Graph Fourier Transform (PhyGraFT), to analyze phylogenetic signals and associations of traits from a similarity network (Fig.1). PhyGraFT was inspired by graph signaling process techniques, such as graph spectral image processing (25). We constructed *k*-nearest neighbor graph from sequence similarity or gene-sharing information, derived graph Laplacian, and performed eigenvalue decomposition. Thereafter, we used graph signaling process framework, graph Fourier transform (GFT) (26), to analyze phylogenetic signals and trait associations from network structure. GFT is the method used to transform graph signals into graph spectrum. The graph is the similarity network, and the graph signal corresponds to phylogenetic trait data. By analyzing the representative graph Fourier coefficients, we can investigate the relationship of evolutionary structures and phylogenetic traits.

**Fig. 1.**
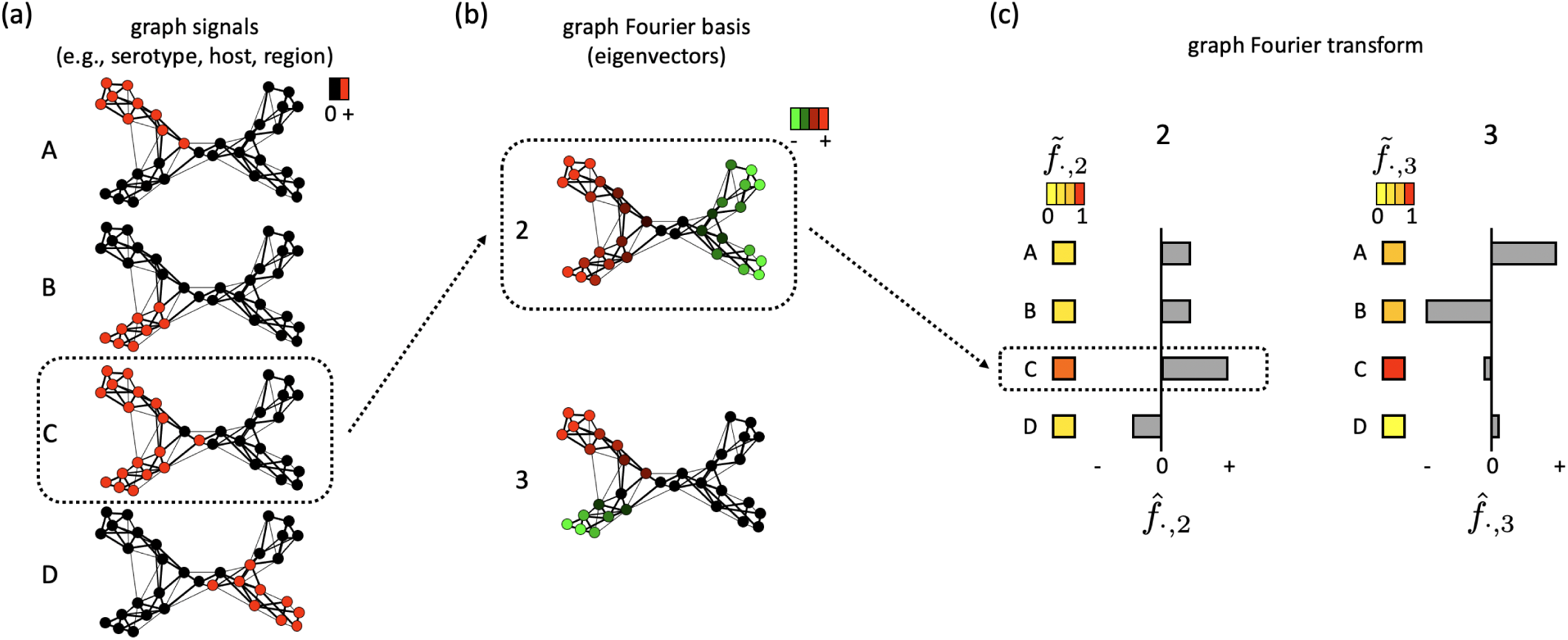
Graphical abstract of PhyGraFT to find phylogeny-trait association structures. (a) Four toy trait data (graph signals) A, B, C, and D, are visualized with the color of nodes, respectively. (b) The second and third eigenvectors (*u*_2_ and *u*_3_), are visualized with the color of nodes, respectively. (c) Visualization of the results of PhyGraFT for *u*_2_ and *u*_3_. For example, the trends of trait A and basis *u*_3_ are consistent, and 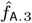 is positively large. The trait C is enriched throughout *u*_3_, and 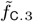 is large.

We firstly validated PhyGraFT, to determine whether it can detect phylogenetic signals from a similarity graph with a simulation dataset. We showed that the relationships of multiple traits could be investigated through graph Fourier coefficients. In addition, we showed that PhyGraFT could detect signals equivalent to existing tree-based methods, even from a similarity graph. These results demonstrated that our network-based approach would be applicable to evolutionary data that cannot be applied with tree-based approach. Thereafter, we applied PhyGraFT for the influenza type A virus datasets and analyzed the phylogenetic signal and associations of traits (serotype, host type, and regions). As a result, we found that GFT effectively identified various network structures, including diversities within the same serotype. We also applied PhyGraFT for the virome genome dataset based on the gene-sharing information, investigated the associations of family, subfamily, and genus data, and found a network structure that contained some genera of different families.

## Materials and Methods

### Outline of the algorithm

PhyGraFT aims to identify phylogenetic signals and associations of various traits from a similarity graph. We constructed graph Laplacian from the similarity graph, calculated the eigenvalues and eigenvectors, and applied a graph signal processing framework to determine phylogeny-trait associations (Fig.1). Thereafter, we explain the detailed procedure of PhyGraFT.

### Graph Laplacian

We represent the weighted adjacency matrix corresponding to the input similarity graph between *n* samples with *A*, and define the symmetrically normalized graph Laplacian matrix *L* by

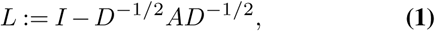

where *I* is the identity matrix, and *D* is the degree matrix.

*L* is a positive semi-definite matrix with one 0 eigenvalue when *A* is a connected graph. Hereafter, we refer *i*-th eigenvalue to *λ*_*i*_ as ascending order (*λ*_1_ = 0 *< λ*_2_ ≤ *λ*_3_ ≤ … ≤ *λ*_*n*_) and corresponding eigenvector to *u*_*i*_. The top eigenvectors, excluding *u*_1_, were used such for graph embedding (20) and phylogenetic tree reconstruction (24).

Since the essential structures of the graph appear in the top eigenvectors, we only used the eigenvalues and eigenvectors up to the top *m*. This procedure reduces the computational time, and allows the application of the algorithm for even a large dataset.

### Graph Fourier Transform

When signals are linked to nodes in a graph (graph signal), the graph Fourier transform, an extension of the standard Fourier transform to graph, is used to transform the graph signal into a graph spectra (26). The basis of graph Fourier transform is the eigenvector of *L*. The graph signal corresponds to trait data, and we denote the signal of trait *i* by *f*_*i*_ ∈ ℝ^*n*^. The graph Fourier transform corresponding to graph signal *f*_*i*_ and eigenvector *u*_*j*_ can be calculated as follows,

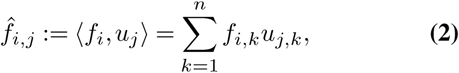

where *f*_*i,k*_ and *u*_*j,k*_ represent *k*-th component of *f*_*i*_ and *u*_*j*_, respectively. In this research, we pre-normalized *f*_*i*_ so that ||*f*_*i*_||_2_ = 1 to compare the graph Fourier coefficients 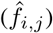 with different traits. With this normalization, 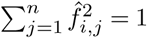 is satisfied.

The values of 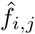 are positively or negatively large if the trait *i* and the eigenvector *u*_*j*_ have a consistent or reverse trend, and almost 0 if they are not relevant (Fig.1). By investigating the 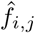 of top eigenvectors, one can determine the evolutionary structures and their associated traits.

We also defined the different indicator 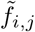 to evaluate the enrichment of trait *i* in the basis *j* regardless of the sign as follows,

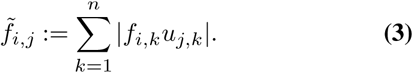

This indicator is helpful in identifying the relationship between trait C and basis 3 in Fig.1, for example.

### Graph Laplacian Regularizer

We developed an indicator to quantify the phylogenetic signal of the trait from the network. The smoothness of the graph signal was evaluated with graph Laplacian regularizer (GLR) (26, 27) as follows,

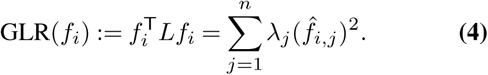

A small GLR(*f*_*i*_) value means that the signal *i* is distributed smoothly on the graph. In other words, the GLR(*f*_*i*_) value will be small when values of the trait in the same clade are similar. This research used GLR to quantify the consistency of trait data and evolutionary network structure. Note that the *f*_*i*_ is pre-normalized so that the mean is 0 and the norm is 1. Because we only calculated top-*m* eigenvalues and eigenvectors, we defined the approximated GLR (aGLR) as follows,

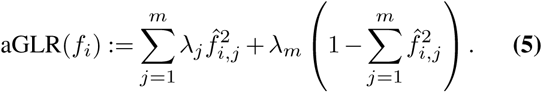

This approximation is based on the fact that 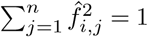 and we use *λ*_*m*_ instead of *λ*_*m*_^*′*^ (*m*^*′*^ *> m*).

### Dataset

#### Construction of k-nearest neighbor graph

In this research, we constructed *k*-nearest neighbor graph (k-NNG) as similarity graph from simulated or real amino acid sequences or gene-sharing information. Thereafter, we used the adjacency matrix of k-NNG to analyze phylogenetic trait data with PhyGraFT.

For amino acid sequence dataset, we used MMSeqs2 Release 13-45111 (28) for all-to-all pairwise sequence alignments and calculated alignment bit score *S*_*bits*_ (*i, j*) between any query sequence *i* and target sequence *j*. The sequence similarity 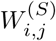 is defined by

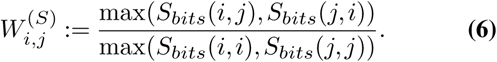

For gene-sharing dataset, we used Jaccard index to calculate the gene-sharing similarity as follows,

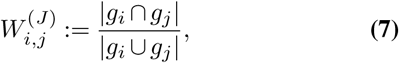

where *g*_*i*_ is the set of orthologous genes in the virome genome *i*.

Thereafter, we defined the adjacency matrix *A* of k-NNG based on the above similarity matrix (*W* ^(*S*)^ or *W* ^(*J*)^). If the two samples *i* and *j* were mutually *k*-nearest neighbors, *A*_*i,j*_ was set to 2, and if the pair was the *k*-nearest neighbor from one side, *A*_*i,j*_ was set to 1, and otherwise *A*_*i,j*_ = 0. We defined the non-zero weight pairs as nearest neighbors of sample *i* when the number of *W*_*i,j*_ *>* 0 was less than *k* for *i*.

#### Simulation Dataset

We used simulation datasets to validate the proposed method by comparing the existing phylogenetic methods. We defined the simulated tree that has four clades as shown in Fig.2(a). Each clade had 25 leaves, and the topology of the tree of each clade was randomly generated with a birth-death stochastic tree. We randomly defined the branch length *α*(1 − ln(*u*(*e* − 1) + 1)) with uniform random value *u. α* is a scaling parameter, and we constructed dataset with *α* = 0.1, 0.25, 0.5, respectively. The branch lengths upstream of each clade were set to *α/*2. Thereafter, we generated approximately 1,000-length simulated amino acid sequences based on WAG model with INDELible v1.03(29) and constructed k-NNG from the sequences (Fig.2(d)). We set *k* = 20 for k-NNG construction and calculated top-20 eigenvalues and eigenvectors.

**Fig. 2.**
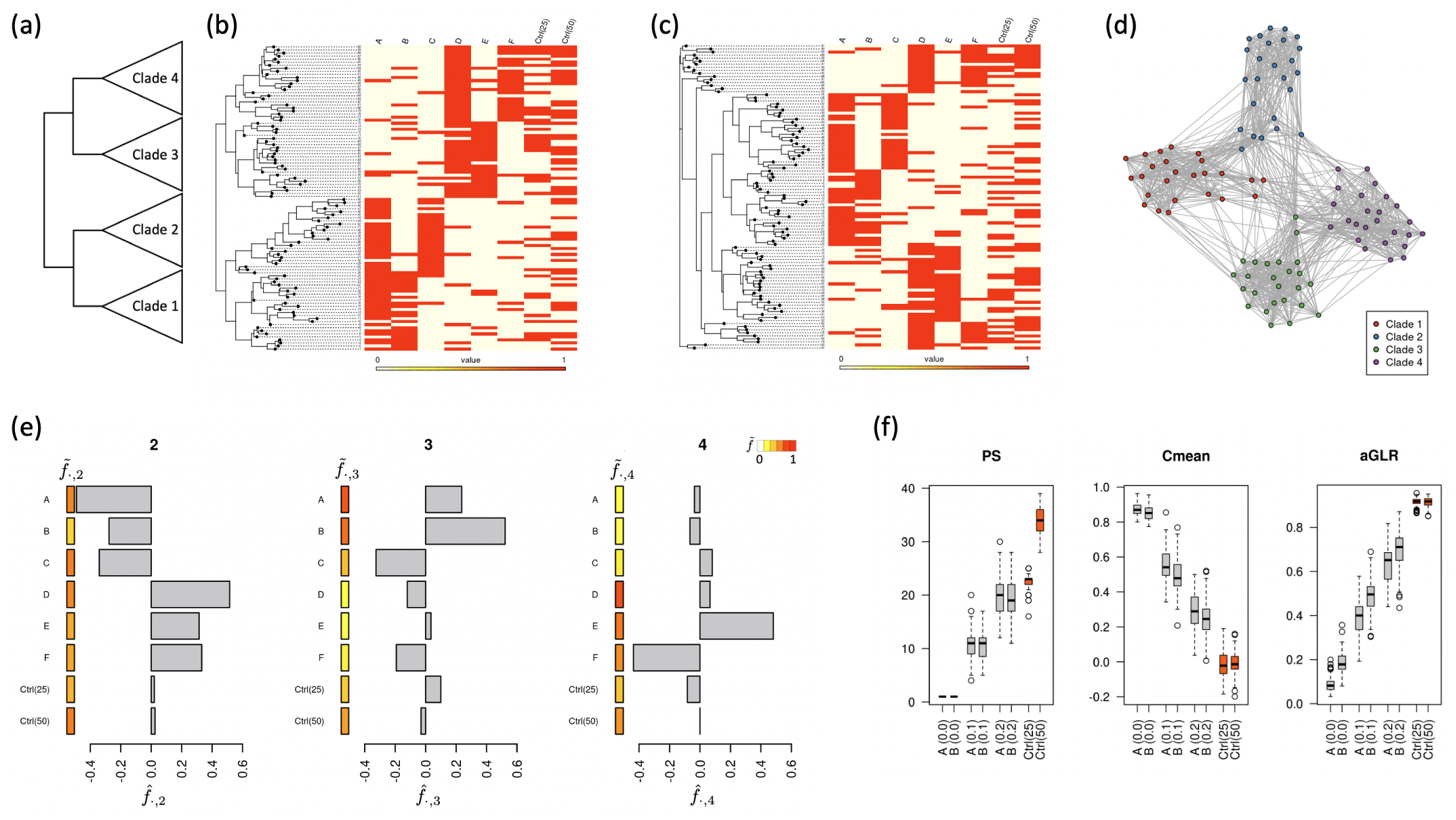
(a) Graphical abstract of simulated tree. (b) Example of simulated tree and trait data with *β* = 0.1. (c) Visualization of reconstructed tree and trait data for the data of (b). (d) Example of k-NNG constructed from simulated amino acid sequences. (e) The graph Fourier coefficients 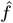and 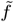 of 2nd, 3rd, and 4th eigenvectors (*u*_2_, *u*_3_, and *u*_4_) for a simulation data. The heat color represents 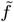 value (red for large 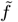 value, and yellow for small 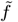 value), and the barplot represents 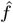 values of each trait data. (f) The PS, *C*_mean_, and aGLR values of positive control traits A and B, and of negative control traits Ctrl(25) and Ctrl(50) for the simulation dataset with *α* = 0.25. A (0.1) means that the values for trait data A with switch error rate *β* = 0.1, for example.

We defined the phylogenetic trait data as binary values. As positive control trait data, we defined six trait data A-F; clade 1-2 had trait A, clade 1 had trait B, clade 2 had trait C, clade 3-4 had trait D, clade 3 had trait E, and clade 4 had trait F (Fig.2(b)). We added the noise to the trait data by switching trait value with probability *β* (*β* = 0, 0.1, 0.2). We also defined negative control trait data, Ctrl(25) and Ctrl(50), by shuffling trait B data and trait A data, respectively (Fig.2(b)). The phylogenetic tree is necessary for tree-based methods. We performed MSA with MAFFT v7.487 (30), calculated the evolutionary distances using dist.ml function in phangorn package (31), and reconstructed the phylogenetic tree using neighbor joining method (32) (Fig.2(c)). We used parsimony score (PS) (3) and *C*_mean_ (33) as the typical tree-based methods to evaluate the trait consistency of the phylogenetic tree. PS and *C*_mean_ were calculated with phangorn package (31) and phylosignal package (6), respectively.

We generated the above simulation dataset of each α (α = 0.1,0.25,0.5) 100 times.

#### Influenza Type A Virus Datasets

We applied Phy-GraFT to the influenza type A virus dataset. We used amino acid sequences of three proteins, hemagglutinin (HA), neuraminidase (NA), and polymerase basic protein 2 (PB2). The sequences were downloaded from the NCBI Influenza virus resource (or database) (https://www.ncbi.nlm.nih.gov/genomes/FLU/Database/nph-select.cgi, downloaded Dec 17, 2021 for HA, Dec 20, 2021 for NA, and Feb 4, 2022 for PB2). We used the sequences with Avian, Swine, or Human hosts, which resulted in 112,965 HA sequences, 95,347 NA sequences, and 69,583 PB2 sequences. Similar sequences were clustered, and the representative sequences of each cluster were obtained using cd-hit v4.8.1 (34, 35). We set the argument “-c 0.98” for envelope proteins (HA and NA) and “-c 0.99” for PB2 for cd-hit usage. After filtering outlier sequences, with significantly smaller similarities than other majority sequences, we obtained 5,106, 4,201, and 2,857 representative sequences for HA, NA, and PB2, respectively. We calculated sequence similarities between representative sequences with MMSeq2 and constructed k-NNGs with *k* = 200 (Fig.3(a), 4(a), and 5(a)). Phylogenetic trait data were also downloaded from the NCBI Influenza virus resource. We obtained the serotype, host, and region information of each sequence. Then, we defined that a cd-hit cluster has a trait if any sequence in the cluster has the trait.

**Fig. 3.**
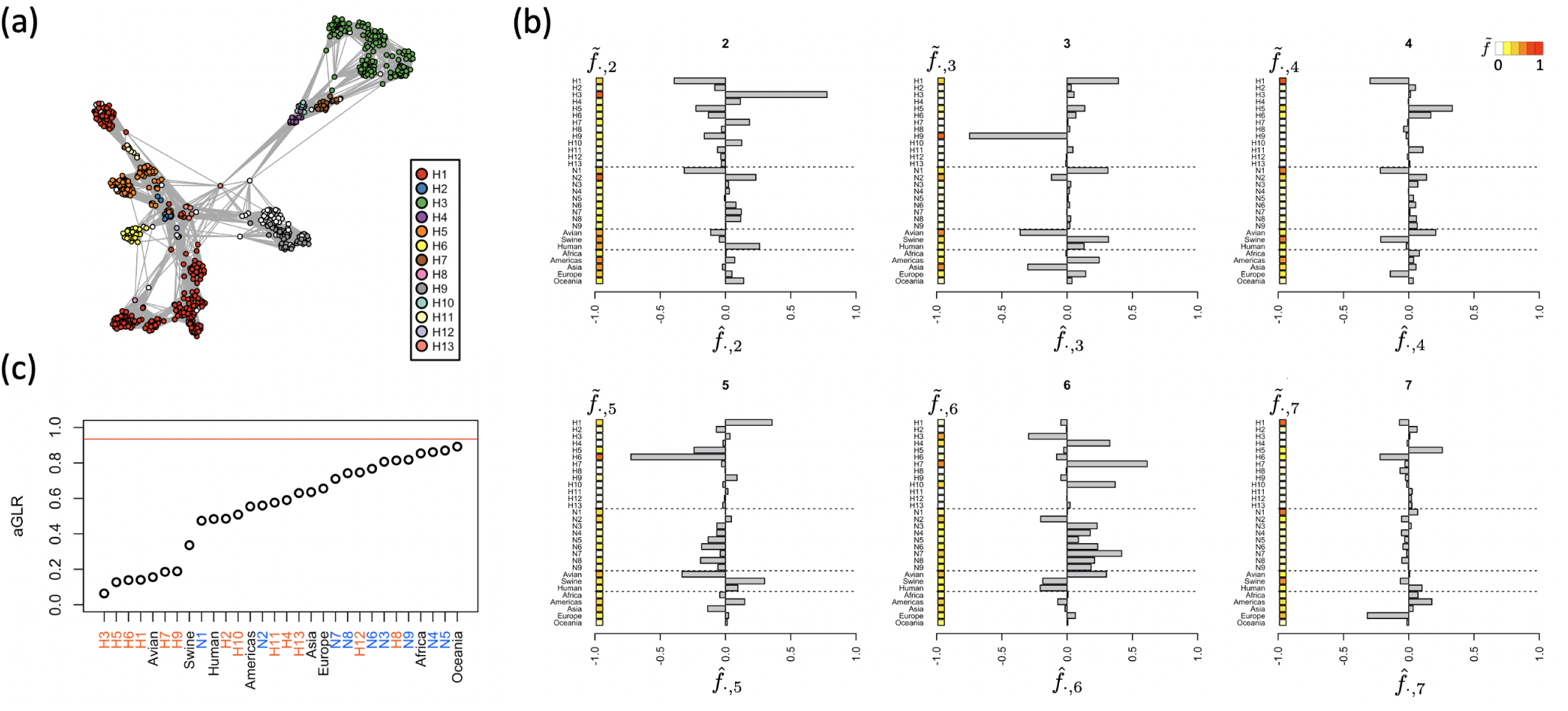
(a) k-NNG constructed from HA sequences of influenza type A virus. For visibility, the networks are randomly down-sampled. Nodes are colored by serotype of HA. (b) The representative 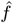 and 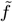 for the HA dataset. The number in the tile represents the index *i* of the eigenvector *u*_*i*_ (*i* = 2 to 7). The heat color represents 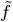 value (red for large 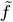 value, and yellow for small 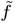 value), and the barplot represents 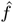 values of each trait data. (c) The aGLR values for the HA dataset. The red line is the averaged aGLR value for negative control trait data that is generated by shuffling randomly selected trait data.

**Fig. 4.**
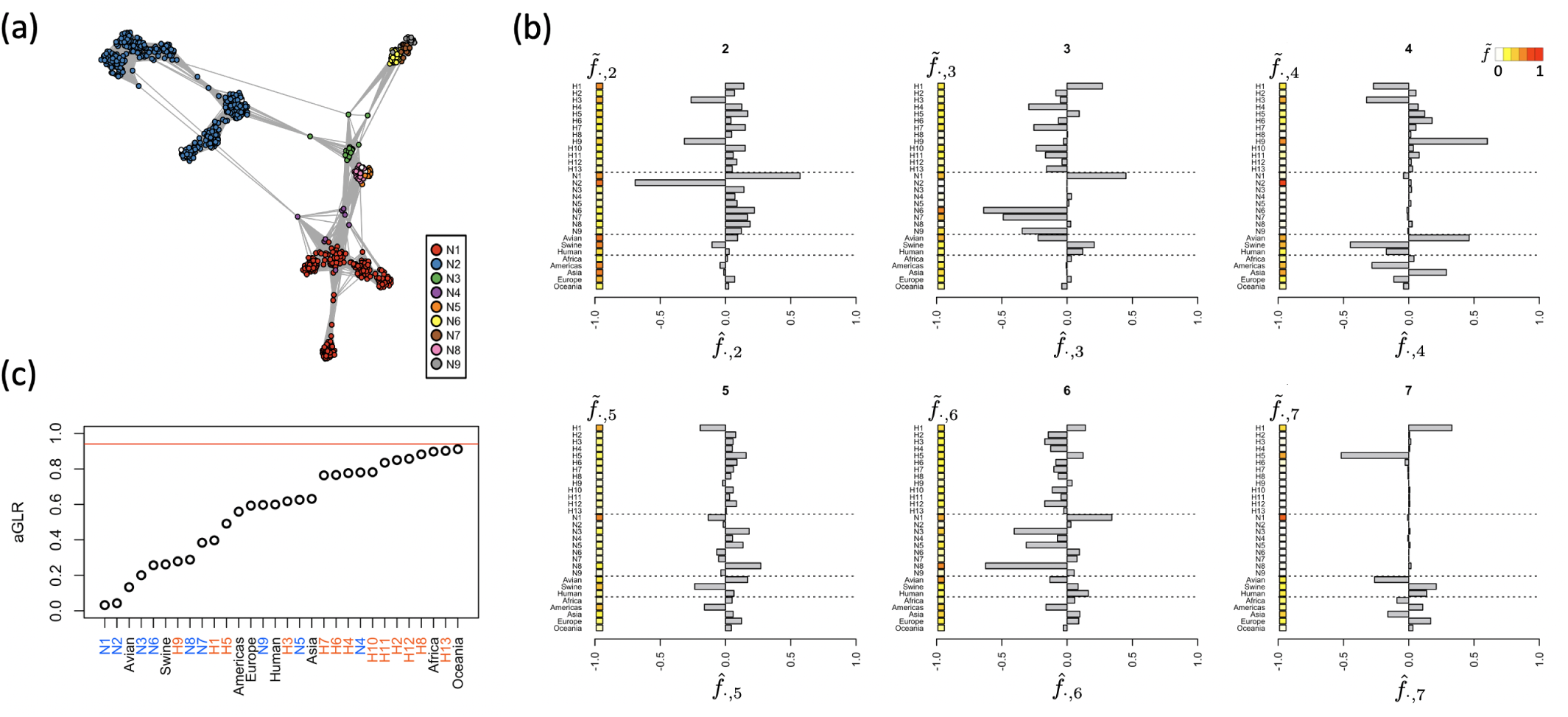
(a) k-NNG constructed from NA sequences of influenza type A virus. For visibility, the networks are randomly down-sampled. Nodes are colored by serotype of NA. (b) The representative 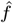 and 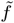 for the NA dataset. (c) The aGLR values for the NA dataset.

**Fig. 5.**
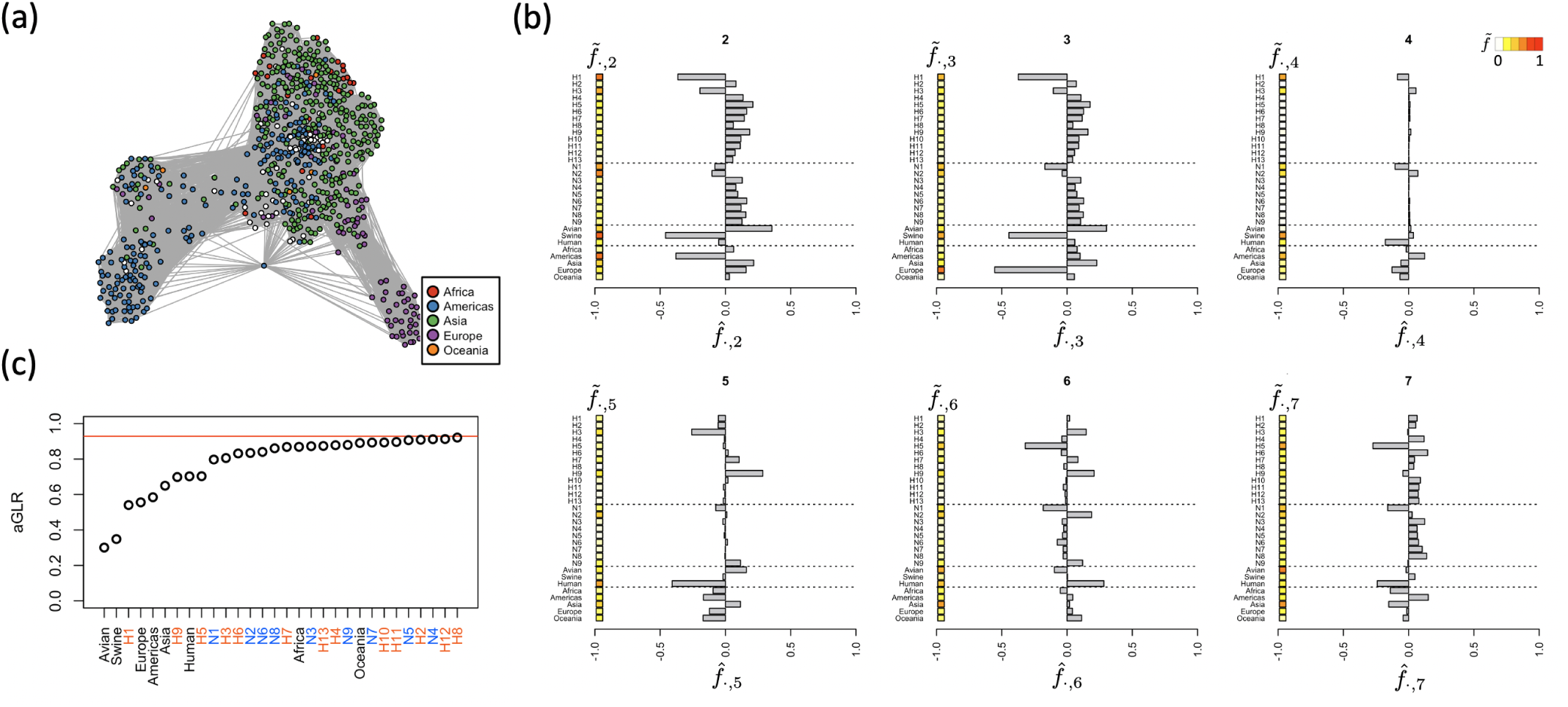
(a) k-NNG constructed from PB2 sequences of influenza type A virus. For visibility, the networks are randomly down-sampled. Nodes are colored by Region. (b) The representative 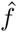 and 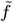 for the PB2 dataset. The number in the tile represents the index *i* of the eigenvector *u*_*i*_ (*i* = 2 to 7). (c) The aGLR values for the PB2 dataset.

#### Virome Genome Gene-Sharing Dataset

We also analyzed the gene-sharing dataset of the Global Ocean Virome genome. The shared protein cluster table of the 2,304 virus genome was downloaded from Supplementary Table 1 of the research of Jang et al. (14). We calculated the similarities between genomes using the Jaccard index and constructed k-NNG with *k* = 100 (Fig.6(a,b)). After filtering the unconnected outlier component of k-NNG, we obtained a 2,301 virus genome network. In this dataset, we regarded the NCBI-family, subfamily, and genus of the virus genome as phylogenetic trait data (downloaded data from Supplementary Table 2 of the research). As there were numerous families, sub-families, and genera, we extracted major trait data only, i.e., family trait data for Myoviridae, Podoviridae, and Siphoviridae, subfamily trait data for 11 subfamilies, such as Autographivirinae and Tevenvirinae, and genus trait data for 12 genera, such as T4virus and L5virus.

**Fig. 6.**
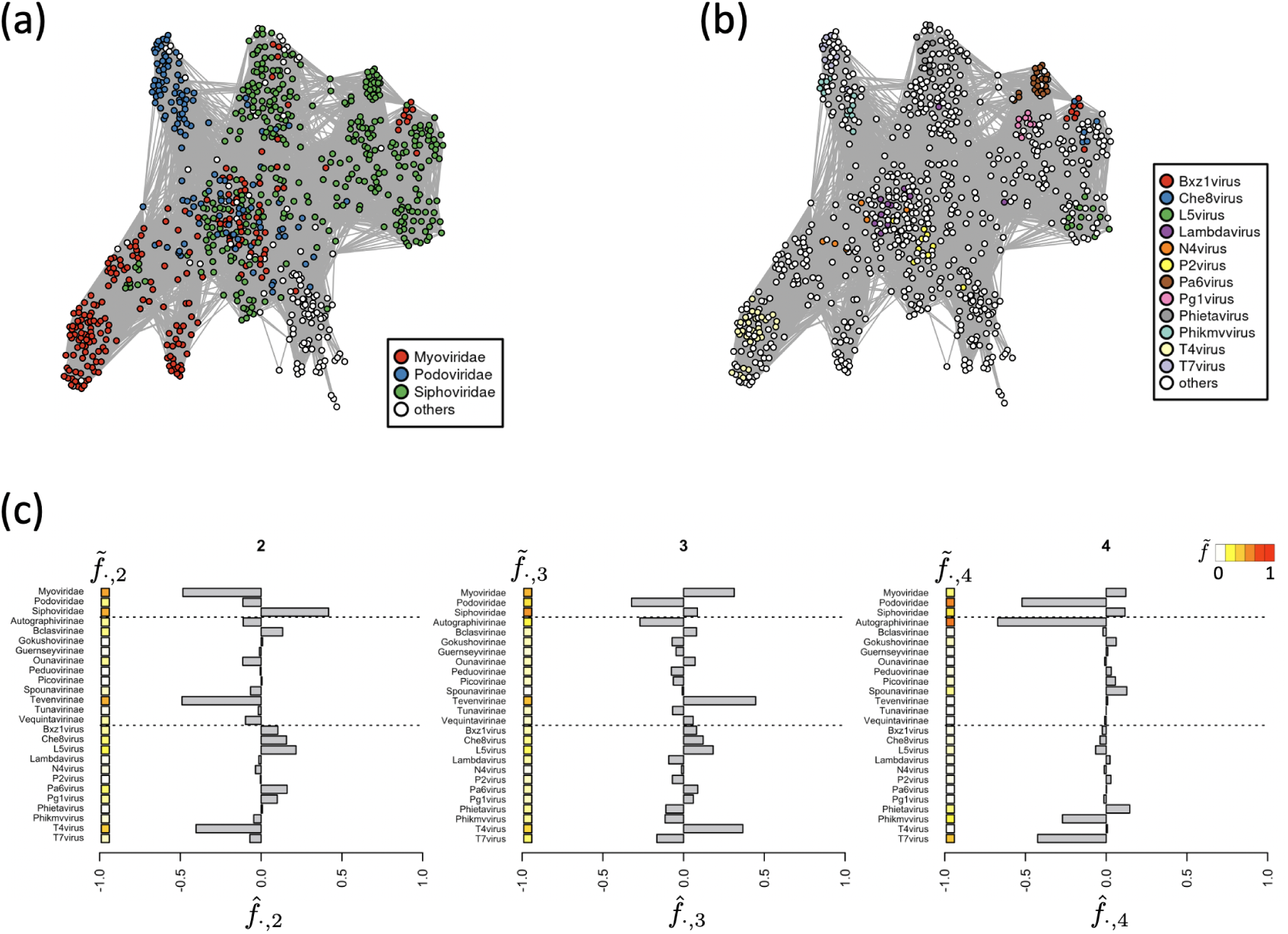
(a,b) k-NNG constructed from virome genome gene-sharing information. For visibility, the networks are randomly down-sampled. Nodes are colored by major (a) NCBI-family and (b) genus, respectively. (c) The representative 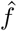 and 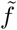 for the virome dataset. The number in the tile represents the index *i* of the eigenvector *u*_*i*_ (*i* = 2 to 4).

## Results

### Validation with the Simulation Dataset

We applied Phy-GraFT for the simulation dataset and evaluate whether PhyGraFT can separate evolutionary structure and find these structure-associated traits. Fig.2(e) shows the representative graph Fourier coefficients 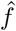 and 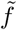 as example data (other examples are shown in Sup. Note 2). 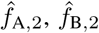, and 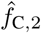 are negatively large, and 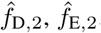, and 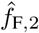 are positively large for the second eigenvector. This result indicates that *u*_2_ represents an evolutionary structure separating clade 1-2 and clade 3-4, and PhyGraFT can find such structure-associated signals. Besides, 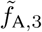 is large, and 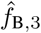 and 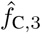 are positively and negatively large, respectively, for the third eigenvector. This result indicates that *u*_3_ separates clades 1 and 2. A similar pattern is observed for clades 3 and 4 for the fourth eigenvector. From these results, the graph Fourier-based method will provide insights into the association of traits and evolutionary structures.

We also validated our indicator aGLR with the simulation dataset by comparing the results with the typical tree-based indicators, parsimony score (PS) and *C*_mean_. PS values, *C*_mean_, and aGLR of positive control (trait A and B) and negative control (Ctrl(25) and Ctrl(50)) for the simulation dataset with *α* = 0.25 are shown in Fig.2(f). The results of the other simulation dataset and the influence of choice of *k* are shown in Sup. Note 1 and Sup. Note 2.

There is a large difference between PS values of positive control with small *β* and negative control Ctrl(25) (*p*-value is 1.8 × 10*−* 117 with Welch’s t-test for A(0.0)), but those of positive control with large *β* are close to those of negative control Ctrl(25) (*p*-value is 3.0 × 10^*−* 8^ for A(0.2)). This is because PS evaluates the trait switch count all over the tree and is not designed to determine the local phylogenetic signal. In contrast, the differences between positive and negative control are clear with both of the values of *C*_mean_ and aGLR even for noisy trait data (*p*-value of *C*_mean_ and aGLR for A(0.2) are 8.0 × 10^*−* 59^ and 2.0 × 10^*−* 59^, respectively). Moreover, these indicators showed almost the same values for Ctrl(25) and Ctrl(50) compared to the PS values. Therefore, *C*_mean_ and aGLR are more useful for evaluating phylogenetic signals. In conclusion, our network-based approach will be applicable for phylogenetic trait analysis even when we only know the similarity network and cannot reconstruct a phylogenetic tree.

### Analysis of the Influenza Datasets

We applied Phy-GraFT for the HA, NA, and PB2 sequence datasets. The representative graph Fourier coefficients and aGLR values are shown in Fig.3, 4, and 5, respectively.

#### HA Dataset

For the HA dataset, various HA types showed large 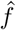 values, such as 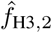 and 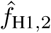 (Fig.3(b)). Moreover, several HA types showed small aGLR values (Fig.3(c)). Therefore, the HA types are significantly associated with the network structures in the k-NNG of HA sequences.

Moreover, the following trait associations are mainly suggested from the values of 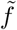 and 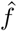 for each eigenvector (Fig.3(b)). *u*_2_ is associated with traits Human/H{3,4,7,10}N{2,6,7,8} and H{1,5,6,9}N1 for positive and negative side, respectively. *u*_3_ is associated with Swine/Americas/H1N1 and Avian/Asia/H9N2 for positive and negative side. *u*_4_ is associated with Swine/H1 throughout and is associated with Swine,Avian/H{1,5,6}N2 and Swine/H1N1 for positive and negative side. *u*_5_ is associated with Swine/H1 and Avian/H{5,6}N{6,8} for positive and negative side. *u*_6_ is associated with Avian/H{4,7,10}N{3,4,6,7,8,9} and Swine,Human/H3N2 for positive and negative side. *u*_7_ is associated with Swine/H1N1 throughout and is associated with Swine/Americas/H{1,5}N1 and Swine/Europe/H{1,6}N1 for positive and negative side.

In particular, 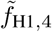 and 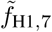 showed large values. This implies that *u*_4_ and *u*_7_ mainly represent the diversities within the H1 sequences. The diversities represented by *u*_4_ and *u*_7_ and its trait associations are visualized in Fig.12(a). As a result, most H1 sequences will be separated into three groups. The first group, the positive side of *u*_4_, is the H1 sequences corresponding to H1N2 strain. The second group, the negative side of *u*_4_ and the positive side of *u*_7_, is the H1 sequences corresponding to Swine,Human/Americas/H1N1 strain. The third group, the negative sides of *u*_4_ and *u*_7_, is the H1 sequences corresponding to Swine/Europe/H1N1 strain. The first group is suggested to be most similar to H5 and H6 sequences, and the second and third groups are relatively similar to H5 and H6, respectively. We also reconstructed the phylogenetic tree using the neighbor joining method and compared the tree with traits (Fig.12(b)). As a result, most H1 sequences are clustered into the clade that rarely contains other HA types. Moreover, the first group of H1 sequences was clustered in a sub-clade of the clade. The H5 and H6 sequences are located apart from the H1 clade on the tree, and the relationship between H1 groups and H5 (or H6) sequences cannot be read off the tree. Such relationships may have been hidden in reconstructing the tree and could be found from the perspective of network-thinking.

In addition, 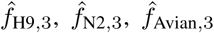, and 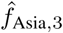 values were negatively large for *u*_3_. This result suggests *u*_3_ will mainly correspond to H9N2 avian strain outbreak in Asia. 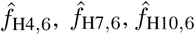, and 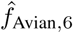 were positively large for *u*_6_, which suggests the existence of the group of avian H4, H7 and H10 sequences.

Thus, PhyGraFT can partition the network into its respective evolutionary structures and evaluate the associated traits.

#### NA Dataset

For the NA dataset, the representative 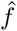 and 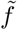 values of various NA types are large (Fig.4(b)), and several NA types showed small aGLR values (Fig.4(c)). Therefore, the NA types are associated with the network structures in the k-NNG of NA sequences.

Moreover, the following trait associations are mainly suggested from the values of 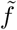 and 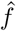 for each eigenvector (Fig.4(b)). *u*_2_ is associated with traits N1 and H{3,9}N2 for positive and negative side. *u*_3_ is associated with Swine/H1N1 and Avian/H{4,7,10}N{6,7,9} for positive and negative side. *u*_4_ is associated with N2 throughout and is associated with Avian/Asia/H9N2 and Swine/Americas/H{1,3}N2 for positive and negative side. *u*_5_ is slightly associated with N1 throughout and is associated with N{1,8} and Swine/H1N1 for positive and negative side. *u*_6_ is associated with N1 and N{3,5,8} for positive and negative side. *u*_7_ is associated with N1 throughout and is associated with H1N1 and H5N1 for positive and negative side.

In particular, 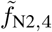 and 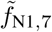 values were large. This means that *u*_4_ will represent the diversity within the N2 sequences, and *u*_7_ will represent the diversity within the N1 sequences. For the fourth eigenvector, 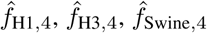, and 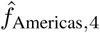 values were negatively large, and 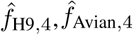 and 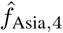 values were positively large. Therefore, there will be two groups within the N2 sequences; one corresponding to the N2 sequences of the H1N2 and H3N2 swine strains from the Americas, and another corresponding to the N2 sequences of the H9N2 avian strain from Asia. For the seventh eigenvector, 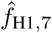 and 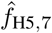 values were positively and negatively large, respectively, which indicates the existence of two groups within N1 sequences; one corresponding to H1N1 strain and another corresponding to H5N1 strain. Moreover, 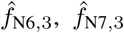, and 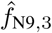 values were negatively large for the third eigenvector, and 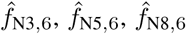 value were negatively large for the sixth eigenvector. These results suggest the evolutionary relevant types of NA can be estimated from the similarity network.

#### PB2 Dataset

For the PB2 dataset, host type and some region traits had small aGLR values (Fig.5(c)). For the second and third eigenvectors, 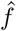 values of H1 and Swine were negatively large. In contrast, 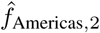 value was negatively large for *u*_2_, and 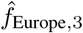 value was negatively large for *u*_3_ (Fig.5(c)). Therefore, both *u*_2_ and *u*_3_ correspond to PB2 sequences of the H1 type swine strains. There are two different groups within these PB2 sequences; one corresponds to the H1 type swine strains outbreak in the Americas, and another corresponds to H1 type swine strains outbreak in Europe. In addition, the sequence groups that correspond to H3N2 and H9N2 human influenza strains or H5N1 avian and human influenza strains were suggested in *u*_6_ and *u*_7_, for example.

### Analysis of the Virome Genome Dataset

The representative graph Fourier coefficients for virome genome dataset are shown in Fig.6(c). For the second eigenvector, 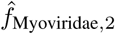 and 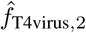 values were negatively large, and 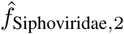 value was positively large. Besides, 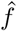 values for some genera, such as 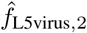 were slightly positively large. These results suggest that the second eigenvector represents the network structure mainly separating Myoviridae, including the T4 virus, and Siphoviridae. Interestingly, although the family of Bxz1virus is Myoviridae, 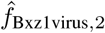 value was slightly positively large. This result suggests that Bxz1virus is more similar to the Siphoviridae groups than the Myoviridae groups in the gene-sharing network (Fig.6(a,b)). For the third eigenvector, 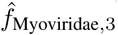 and 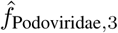 values were positively and negatively larege, and this result suggest that the third eigenvector represents the network structure mainly separating Myoviridae and Podoviridae. For the fourth eigenvector, 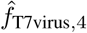 and 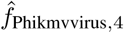 values were negatively large. Phikmvvirus is a T7-like virus, and therefore, such genus relationships can be found from the network-based approach.

## Discussion

In this research, we developed a novel network-thinking approach PhyGraFT to identify evolutionary signals and associations of traits based on the graph signaling process framework. Several graph Laplacian-based methods have been proposed for evolutionary studies (21–24), and this research is a further development of these approaches by introducing the GFT framework into phylogenetic analysis. We applied PhyGraFT for influenza type A virus datasets and virome gene-sharing dataset and demonstrated that we could locate various evolutionary structures and these associated traits from similarity networks. PhyGraFT provides a flexible network-based analysis method and is useful for further development of network-thinking approaches. The source code of PhyGraFT and the processed datasets are available at https://github.com/hmatsu1226/PhyGraFT.

In the Artificial Intelligence area, research on graph neural networks (GNN) has recently advanced (36). The graph Laplacian and GFT are closely related to the GNN (37). Therefore, the network-based phylogenetic analysis approach has further development potential by introducing findings in those areas.

Recently, GFT has also been used for several biological analyses, including single-cell RNA-sequencing data analysis (38). Our research demonstrated the effectiveness of GFT for phylogenetic network signal analysis, and will promote network-thinking for various biological research, such as bacterial evolution, including horizontal gene transfer (17) and antibody evolution (18).

## ACKNOWLEDGEMENTS

This work was supported by the Japan Society for the Promotion of Science (Grant number 21K17858 to H.M., JP19H05679 and 19H05688 to M.M., JP20H05582 and JP22H04891 to Tsukasa Fukunaga). We thank Tsukasa Fukunaga, Momoko Hayamizu, and Masao Ueki for helpful discussion about the algorithm and Wataru Iwasaki for advice to improve the article.

---

**Supplementary Note 1: Validation with the Simulation Dataset**

The results for the simulation dataset with *α* = 0.1 and 0.5 are shown in Fig.7. Both of *C*_mean_ and aGLR can distinguish positive and negative control trait data as described in the main text even with large *α* and large *β*.

**Fig. 7.**
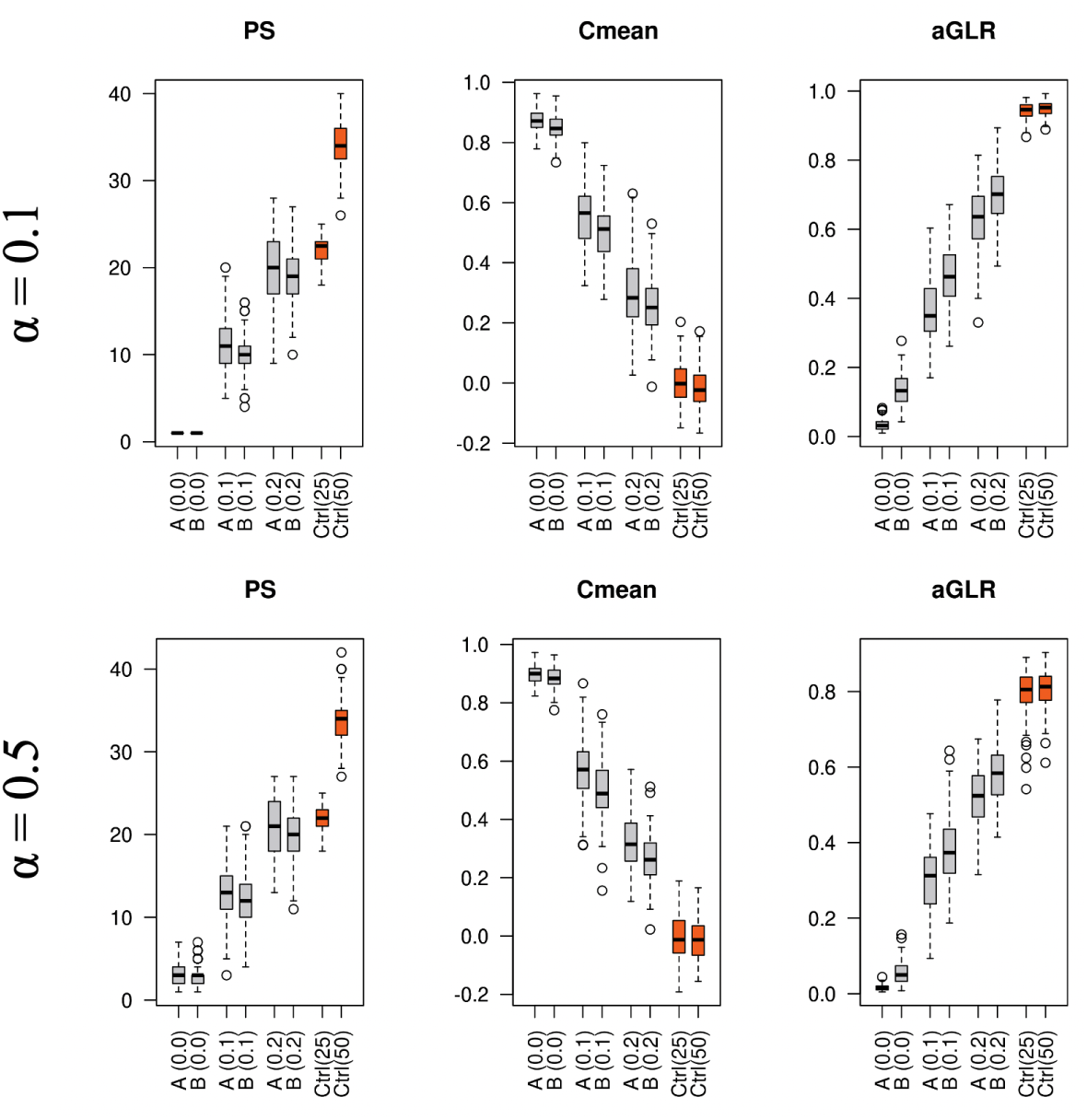
PS, *C*mean, and aGLR values of positive control traits A and B, and of negative control traits Ctrl(25) and Ctrl(50) for the simulation dataset with *α* = 0.1 and 0.5.

We also show the results of the representative 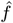 and 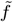 values of PhyGraFT for some simulation data (*α* = 0.25) in Fig.8. In most cases, the clades 1-2 and 3-4 are separated in the second eigenvectors. The relationships of clade 1 and 2 or clade 3 and 4 appeared in other top eigenvectors. For some cases, these two separations appeared in the same eigenvector.

**Fig. 8.**
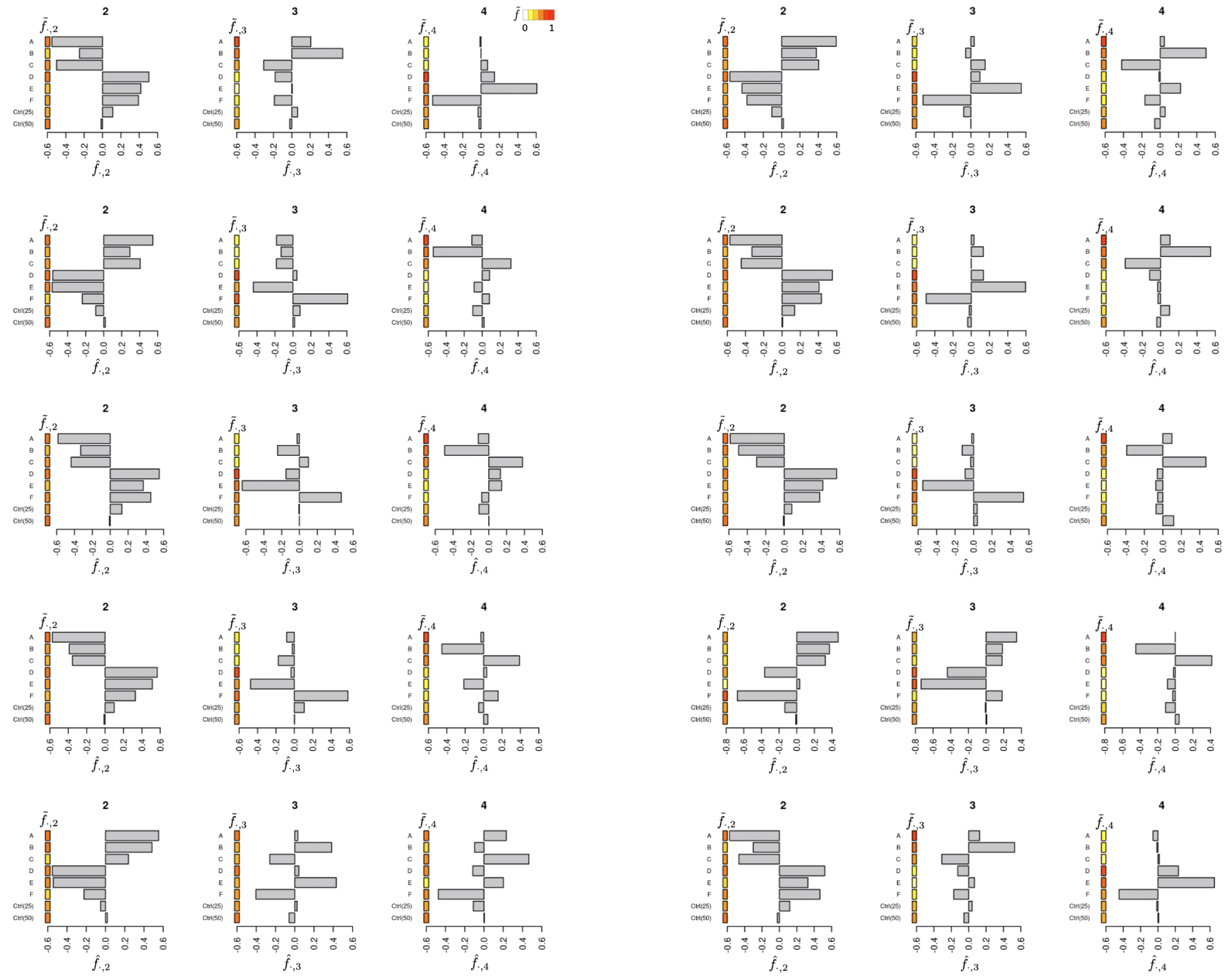
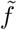 and 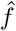 for the second, third, and fourth eigenvectors of 10 example simulation datasets (*α* = 0.25).

**Supplementary Note 2: Analysis of the choice of** *k* **in k-NNG**

We evaluated the influence of the choice of *k* in k-NNG for PhyGraFT with the simulation dataset (*α* = 0.25). The example k-NNG with *k* = 5, 10, 20, 40, and 80 are shown in Fig.9.

**Fig. 9.**
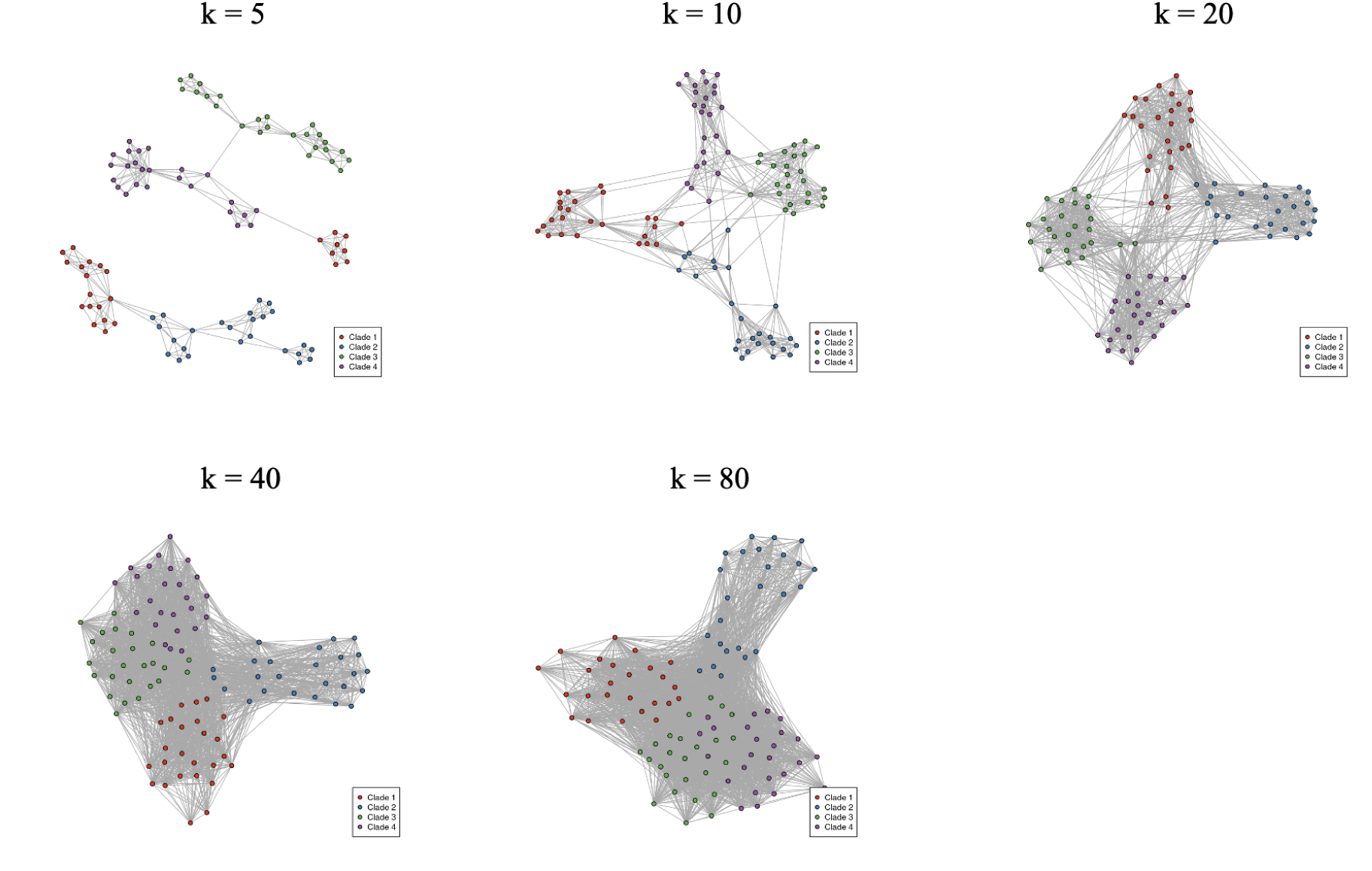
Example k-NNGs with *k* = 5, 10, 20, 40, and 80.

The results of aGLR are shown in Fig.10, and the example representative 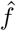 and 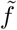 are shown in Fig.11. Although the differences in aGLR values between positive and negative control are clear (Fig.10), some relationships between traits become unclear with *k* = 5 (Fig.11). This must be because the k-NNGs become an unconnected graph, and the global structure information of the original evolutionary structures is diminished in many cases. On the other hand, the differences in aGLR values between positive and negative control are less clear with large *k* (Fig.11). As the fully connected graph has no trait information, highly dense k-NNG is unsuitable for PhyGraFT. The differences in aGLR between positive and negative control are clear, and we can determine the trait associations based on 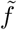 and 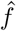 in most cases with *k* = 10, 20, and 40. Therefore, it is preferable to choose *k*, such that k-NNG is the connected graph and is yet sparse.

**Fig. 10.**
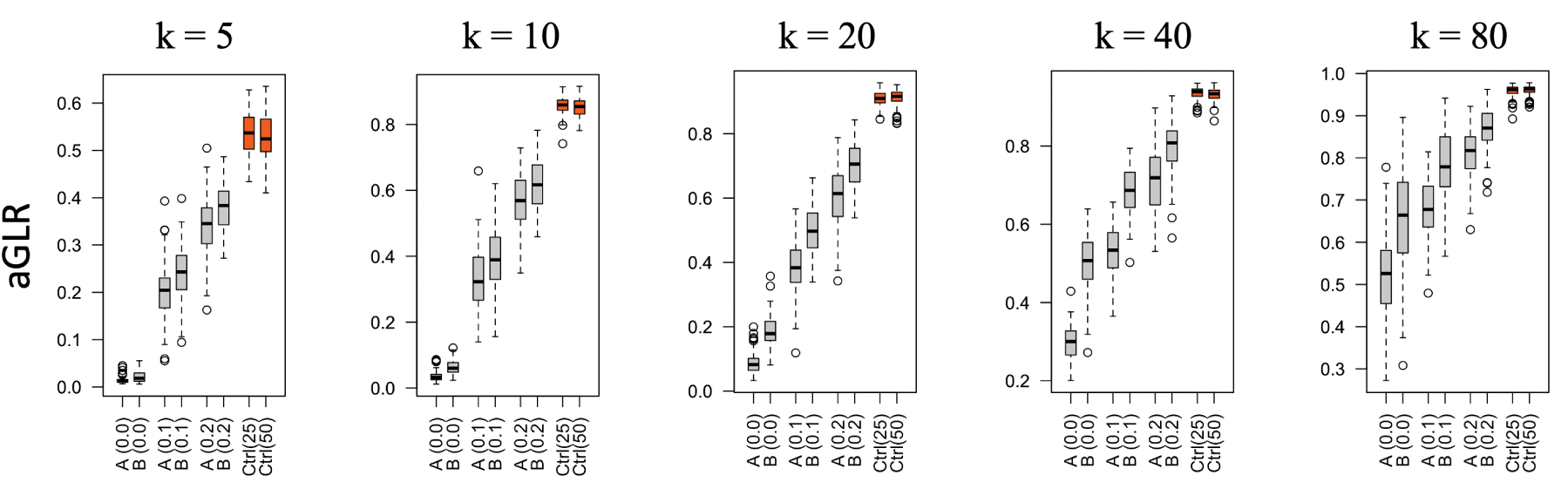
aGLR values of positive and negative control traits for k-NNGs with *k* = 5, 10, 20, 40, and 80.

**Fig. 11.**
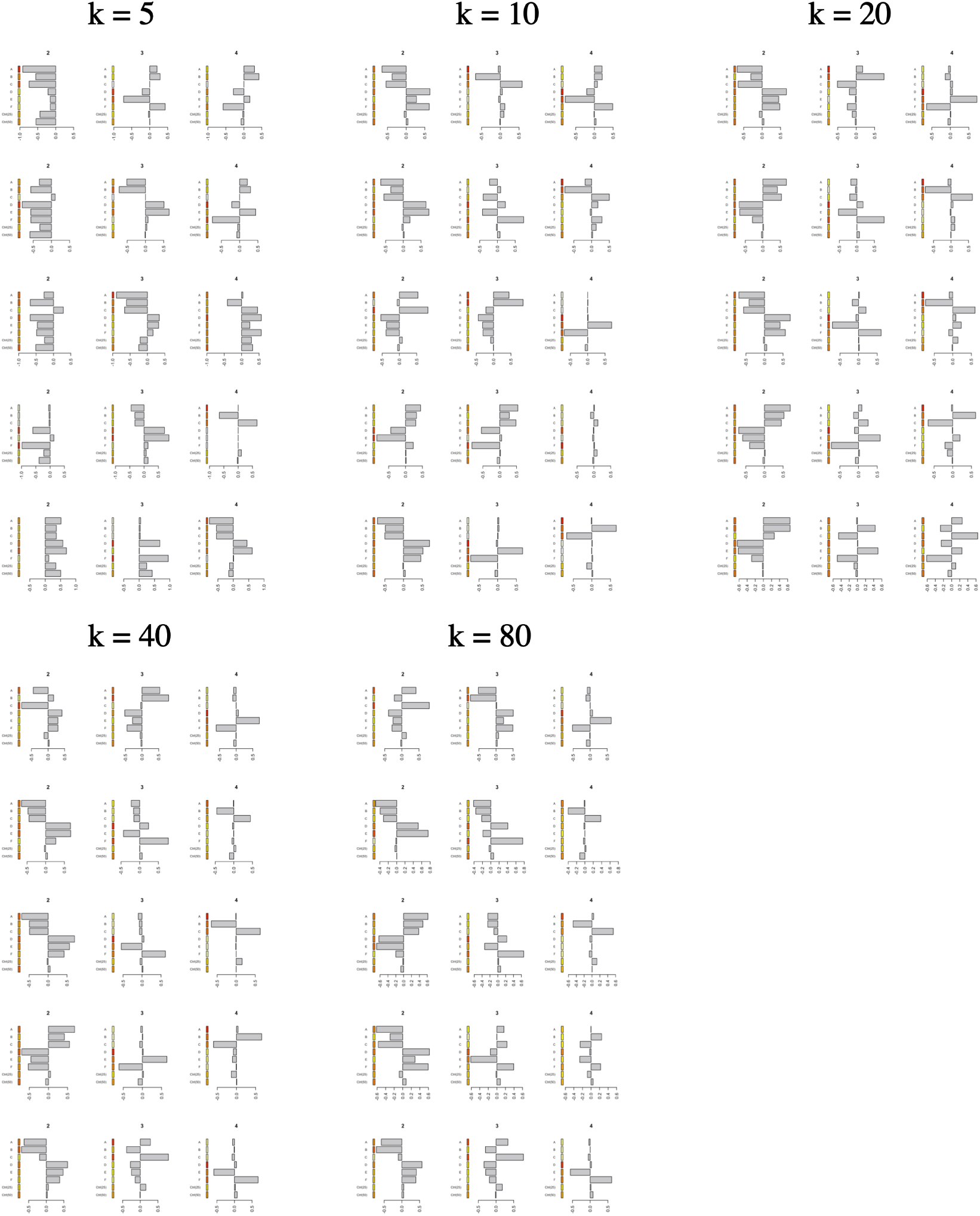
Top 3 representative graph Fourier coefficients for 5 example simulation datasets with *k* = 5, 10, 20, 40, and 80.

**Supplementary Note 3: Additional analysis of the HA dataset**

As mentioned in the main text, *u*_4_ and *u*_7_ represent the diversities of H1 sequences. Fig.12(a) show the scatter plot of *u*_4_ and *u*_7_ colored by each trait. As a result, H1 sequences are suggested to be separated into three groups; the first group corresponds to the H1N2 strain, the second group corresponds to the Swine,Human/Americas/H1N1 strain, and the third group is the Swine/Europe/H1N1 strain. The first group is suggested to be most similar to H5 and H6 sequences.

**Fig. 12.**
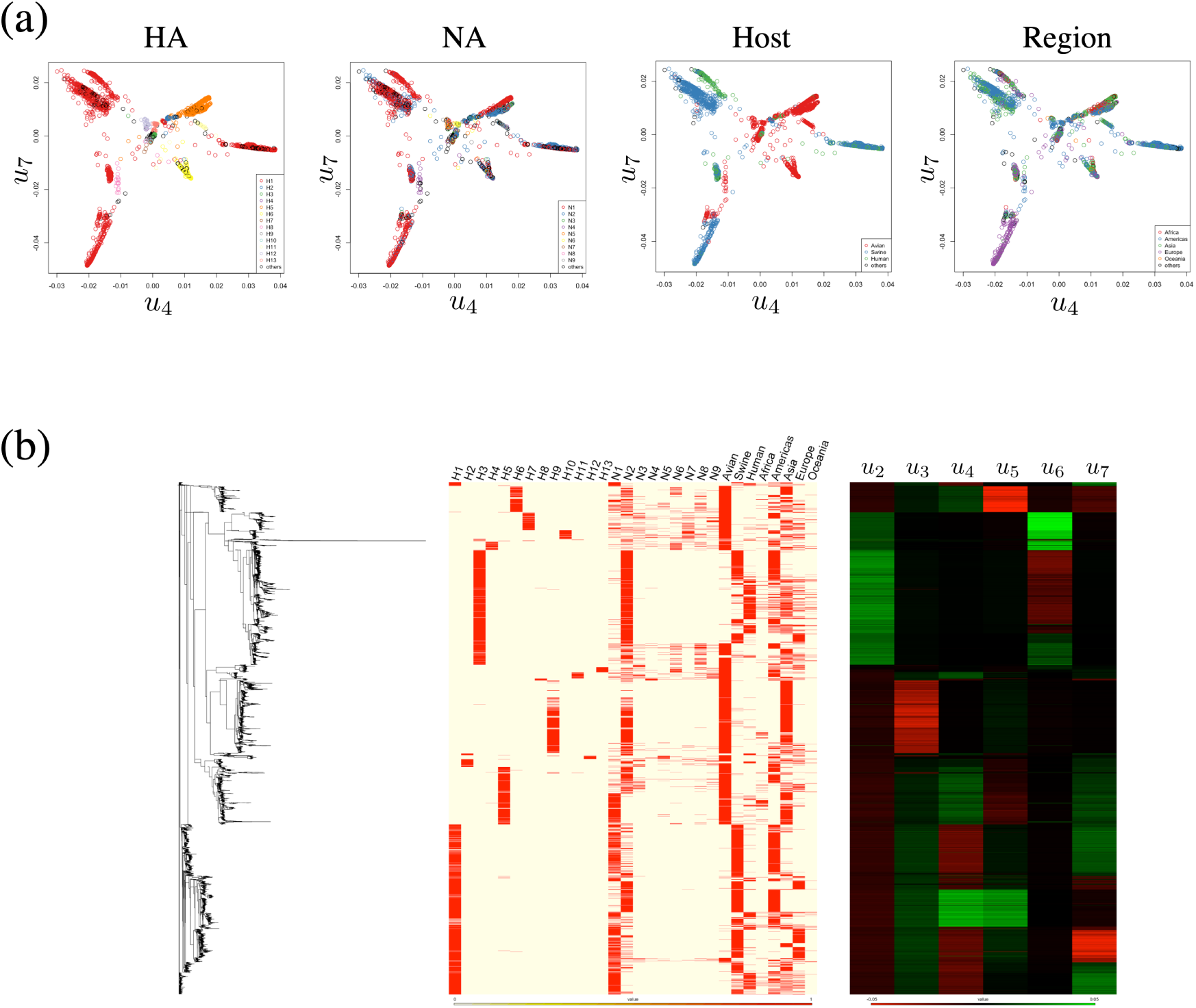
(a) The scatter plot of *u*_4_ and *u*_7_ colored by HA type, NA type, Host type, and Region. (b) The reconstructed tree with neighbor joining method and the heatmaps of corresponding trait data and eigenvectors (*u*_2_-*u*_7_).

We also reconstructed the phylogenetic tree with the neighbor joining method and compared the tree with traits and the eigenvectors (Fig.12(b)). For tree reconstruction, we performed multiple sequence alignment with MAFFT v7.487 (30) and calculated the evolutionary distances using dist.ml function and reconstructed the tree using NJ function in phangorn package (31). A small fraction of the HA sequences contained amino acid code J, indicating Leucine or Isoleucine. We converted J to I (Isoleucine) for evolutionary distance calculation.

Most H1 sequences are clustered in a clade, and the first group (H1N2) of H1 sequences was further clustered in a sub-clade of the clade. The H5 and H6 sequences are located apart from H1 sequences on the tree, and the relationship between H1 groups and H5 (or H6) sequences cannot be read off.

